# Discovery of a potent anti-Zika virus benzamide series targeting the viral protein NS4B

**DOI:** 10.1101/2025.10.09.681361

**Authors:** Donghoon Chung, Yuka Otsuka, Eunjung Kim, Sultan Ullah, Nicole M. Kennedy, Khac Huy Ngo, Jinjoo Kang, Reece Tomlison, Brian Alejandro, Koji Barnaby, Jeffery Miller, Justin Shumate, Chwee Fang Teo, Yaw Bia Tan, Che Shin Chew, CongBao Kang, Dahai Luo, Louis Scampavia, Timothy P. Spicer, Thomas D. Bannister

## Abstract

1.

Zika virus (ZIKV), a member of the *Flaviviridae* family, causes significant public health concerns through congenital Zika syndrome and Guillain-Barré syndrome, yet no effective anti-ZIKV drugs or vaccines are available. To address this critical need, we conducted phenotypic, cytopathic effect-based, high-throughput screening followed by medicinal chemistry optimization and discovered novel benzamide anti-ZIKV leads. Current best compounds demonstrated superior potency (EC_50_ values 40-400 nM, CC_50_ > 50 µM) compared to NITD-008, the most potent known anti-ZIKV agent. Time-of-addition assays, resistant virus selection studies, and biophysical binding experiments confirmed that NS4B interference constitutes the primary antiviral mechanism. Notably, resistance mutations mapped to the C-terminus of NS4B, distinct from other flavivirus NS4B inhibitors targeting dengue or yellow fever viruses, revealing novel insights into a critical function of the region. These findings establish NS4B as an Achilles’ heel for flaviviruses and support the development of pan-flavivirus therapeutics targeting this essential viral protein.

**AUTHOR SUMMARY:** Since its pandemic spread in 2015-2016, Zika virus infection remains a significant public health threat worldwide. The virus can cause severe brain damage in developing babies and serious neurological complications like Guillain-Barré syndrome in adults. Despite these devastating consequences, we currently lack effective medicines or vaccines to prevent the virus from spreading through communities or from mothers to their unborn children. To address this critical gap, we conducted a large-scale screening of chemical compounds and discovered a promising new class of molecules that can effectively stop Zika virus from replication. Using medicinal chemistry techniques, we were able to make these compounds even more potent against the virus.

In follow-up studies, we found that our compounds work by interfering with a specific viral protein called NS4B, which the virus needs to replicate its genome within the cell. Remarkably, other research teams studying related other flaviviruses (e.g., dengue and yellow fever virus) have independently discovered that this same protein is a vulnerable target. Our findings suggest that NS4B represents a universal weakness across the entire flavivirus family, making it an attractive target for developing broad-spectrum antiviral treatments.

## 3. INTRODUCTION

Zika virus (ZIKV) and other flaviviruses are important emerging pathogens of high pandemic potential. Most of these viruses use mosquitoes as a vector and are globally endemic in tropical and subtropical areas(1). Due to climate change, the habitats of carrier mosquito species (i.e., *Aedes albopictus* and *Aedes aegypti*) are rapidly expanding along with regions of susceptibility to outbreaks and local epidemics of these diseases(2). ZIKV was first isolated in Africa in 1947, spread to the Americas in 2015, and has been continuously circulating in many countries in South America. In 2024, the Pan American Health Organization (PAHO) confirmed more than 44,000 cases in the Americas, with >90% of cases in Brazil(3). As of 2024, Zika infections had been reported in 92 countries, however, posing a threat to ∼60 additional countries based on the expanded range of the carrier mosquitoes(4), indicating a growing risk for multi-continent outbreaks.

ZIKV infection typically results in self-limiting febrile clinical outcomes(5). However, 5–15% of infected pregnant women exposed to ZIKV have babies with Zika-related birth defects including infant microcephaly and other cognitive developmental defects, known as congenital Zika syndrome. In adults, Guillain-Barre syndrome, stillbirth, and miscarriages have been reported(6–8). ZIKV can spread via a human or urban cycle, in which infected humans drive its spread(9). ZIKV urban epidemics occur through the transmission of infectious virus in the blood to naïve individuals via peridomestic/domestic mosquitoes (*Aedes spp.)* or through contact with bodily fluids (e.g., semen, vaginal fluid, saliva, urine, breast milk). Additionally, ZIKV can lead to a persistent infection, producing virus in bodily fluids that clears only after many months or even up to a year in certain cases(10–12).

ZIKV persistence in semen allows sexual transmission, passing to the mother and to a fetus, with resultant congenital Zika syndrome. Therefore, it is reasonably expected that early anti-ZIKV treatment which directly reduces the viral burden in the blood or bodily fluid of infected people will quickly minimize local viral spread, reducing pandemic potential. A treatment reducing this persistence could not only curb transmission but reduce fetal effects of Zika syndrome, such as microcephaly. However, no antivirals nor vaccines for ZIKV infection have been approved, emphasizing a pressing need for such agents.

Phenotypic high-throughput based screening (HTS) approaches have often led to the discovery of novel antiviral compounds with new antiviral mechanisms of action, especially where no established targets are had been identified(13–15). Prior phenotypic screens for anti-flaviviral agents identified small molecules targeting DENV (e.g., JNJ-A07, JNJ-1802 and NITD-688), and YFV (BDAA series)(16–19), several of which have demonstrated efficacy in preclinical studies and entered clinical development(20,21). Interestingly, mechanistic studies established that the antiviral targets of each of these anti-DENV compounds is the viral membrane protein NS4B, suggesting that targeting NS4B might also have utility for other flaviviruses. None of these molecules demonstrated anti-ZIKV activity, however, and no effective anti-ZIKV compounds are known thus far, though the nucleoside analog NITD-008, a viral polymerase inhibitor, has pan-flavivirus activity, including ZIKV (22).

Here, we present a novel benzamide lead series discovered from a phenotypic HTS for small molecules that inhibit ZIKV replication. Our mechanism of action studies showed that the series selectively targets ZIKV by binding to and disrupting the function of ZIKV NS4B. Initial screening hits with modest potency were optimized in a medicinal chemistry campaign, improving anti-ZIKV EC_50_ values to below 100 nM with greater than 1000-fold virus yield reduction following exposure to optimized compounds at 1 µM in cell-based assays. Our study establishes that the known importance of NS4B in DENV and YFV as a valid antiviral target extends to ZIKV. This suggests a similar key role for NS4B in other important members of the flavivirus family (e.g., WNV, JEV, and TBEV), suggesting that opportunistically screening or optimizing currently available anti-NS4B compounds to target these other flaviviruses, for which limited, or no treatment options are available, may be a viable strategy for anti-flaviviral drug development.

## 4. RESULTS

### 1. Identification of anti-ZIKV hits from a CPE-based HTS campaign

We previously developed a phenotypic, CPE-based HTS assay for antiviral discovery for ZIKV(23). To avoid finding host-targeting pyrimidine synthesis inhibitors, which have been often identified as poorly tractable hits in other cell-based phenotypic antiviral HTS campaigns(14,24–27), the assay media was supplemented with 50 µM of uridine. Uridine supplementation effectively subverted the antiviral effects of brequinar, a cellular dihydroorotate dehydrogenase inhibitor, in our assay (See **Fig. S1**). For executing an automated HTS campaign to discover new compounds with anti-ZIKV activity, the assay was successfully miniaturized to a 1536-well plate format. The assay performance was robust with the overall Zʹ > 0.8, and the antiviral activity of NITD-008 (EC_50_ = 0.8 ± 0.037 µM, n=10) agreed with previously published data(22).

The UF Scripps diversity small molecule library of 650K compounds was screened at a single concentration in the assay (See **Fig. S2**). Active “cherry pick” compounds were retested for reproducible activity at a higher cutoff, while druglikeness filters (e.g., removing high M.W. compounds, PAINS, and frequent assay hitters), removed some hits from follow-up consideration. 91 compounds of interest remained and were repurchased from vendors and retested (along with selected purchased structural analogs) using several lines of independent assays, including a triage of two orthogonal secondary concentration-dependent assays (a CPE-readout and a reporter-based ZIKV assay), and finally a CPE-based anti-CHIKV counter assay. These efforts gave a small set of ∼25 compounds of interest with activity in one or both ZIKV assay formats, no activity vs. CHIKV (an unrelated alphavirus), and low cellular cytotoxicity suggestive of valid antiviral activity. We found several hits sharing a common benzamide core having significant antiviral activity in the two orthogonal assays (EC_50_ < 2 µM, **Table 1**), activity that was not compound batch dependent.

**Table 1.**
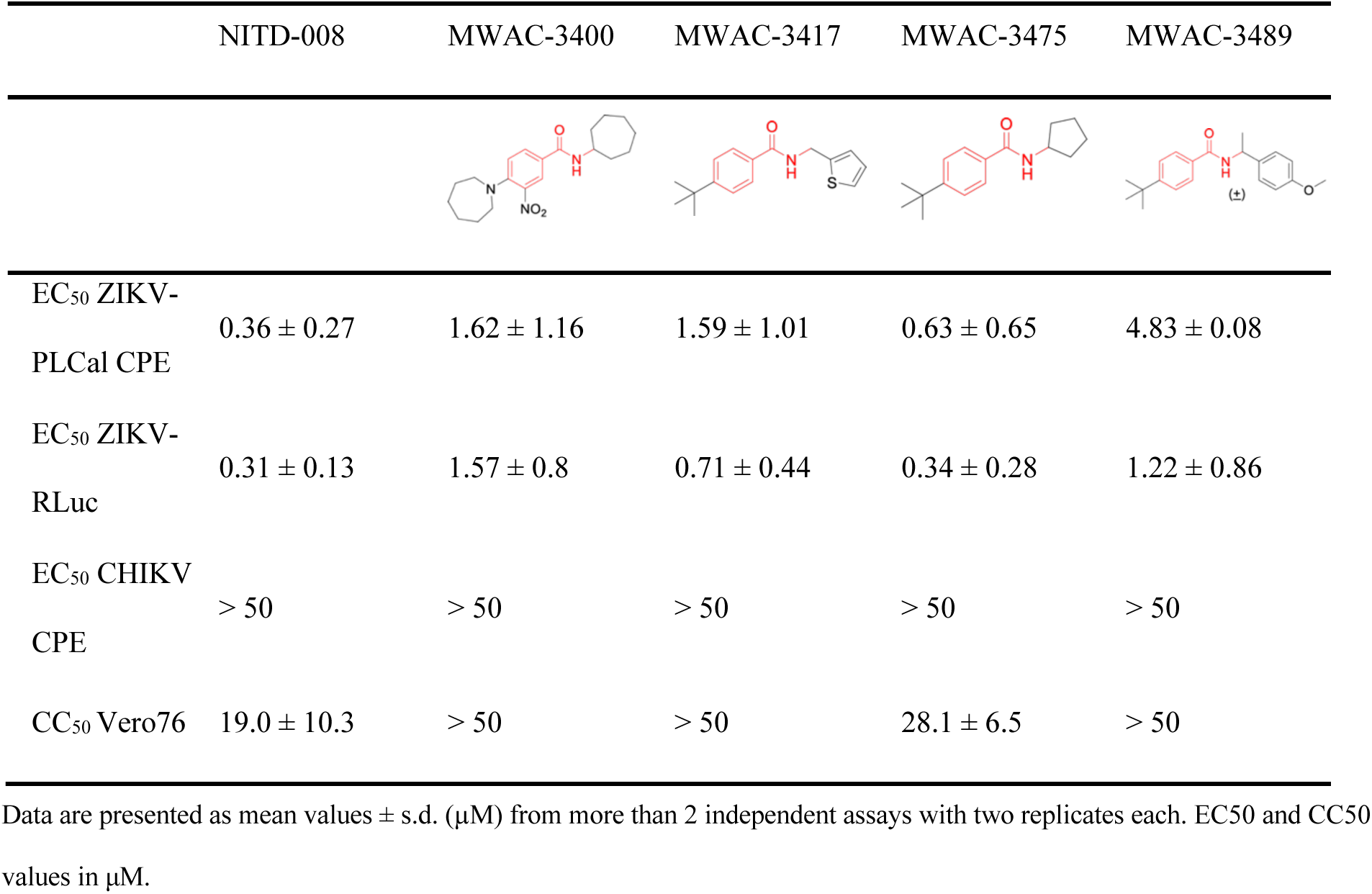
Antiviral activity against ZIKV and CHIKV, and cytotoxicity of the benzamide series hits.

|  | NITD-008 | MWAC-3400 | MWAC-3417 | MWAC-3475 | MWAC-3489 |
| --- | --- | --- | --- | --- | --- |
| $EC_{50}$ ZIKV-PLC<sub>al</sub> CPE | $0.36 \pm 0.27$ | $1.62 \pm 1.16$ | $1.59 \pm 1.01$ | $0.63 \pm 0.65$ | $4.83 \pm 0.08$ |
| $EC_{50}$ ZIKV-RLuc | $0.31 \pm 0.13$ | $1.57 \pm 0.8$ | $0.71 \pm 0.44$ | $0.34 \pm 0.28$ | $1.22 \pm 0.86$ |
| $EC_{50}$ CHIKV CPE | > 50 | > 50 | > 50 | > 50 | > 50 |
| CC<sub>50</sub> Vero76 | $19.0 \pm 10.3$ | > 50 | > 50 | $28.1 \pm 6.5$ | > 50 |

Data are presented as mean values ± s.d. ( $\mu M$ ) from more than 2 independent assays with two replicates each. $EC_{50}$ and $CC_{50}$ values in $\mu M$ .

**Table 1** compares data for top hits, the most potent being the low molecular weight (245) benzamide MWAC-3475, which showed EC_50_ values comparable to NITD-008, a nucleoside analog with a broad-spectrum anti-flavivirus activity. All compounds had no observed activity against CHIKV, suggestive of potential specificity for ZIKV.

### 2. Assessment of ZIKV-specific antiviral activity

To validate the observed anti-ZIKV activity and to understand the mechanism of action (MOA) of the benzamide series (termed the MWAC-3475 series hereafter), we first sought to evaluate its antiviral activity against various strains of ZIKV and against other flaviviruses, including DENV, Yellow fever virus (YFV), Japanese encephalitis virus (JEV), and West Nile virus (WNV). MWAC-3475 series compounds showed antiviral activities for all the ZIKV strains tested with an EC_50_ range of 0.67-1.76 µM but showed no measurable antiviral activity for other flaviviruses up to 50 µM (**Table 2**). This differs from NITD-008, which demonstrated a broad-spectrum antiviral activity for all tested flaviviruses with a low micromolar EC_50_ values. This result showed that MWAC-3475 series is a ZIKV-specific antiviral, potentially targeting a viral motif unique to ZIKV.

**Table 2.** Anti-flavivirus activity of the MWAC-3475 series.

| Compounds and EC<sub>50</sub> values (μM) | | | | | |
| --- | --- | --- | --- | --- | --- |
| Tested virus/strain |  | NITD-008 | MWAC-3400 | MWAC-3417 | MWAC-3475 |
| ZIKV | MR766 | 1.76 ± 0.2 | 9.18 ± 1.07 | 9.95 ± 0.61 | 5.93 ± 1.26 |
| | DAK-41525-MA | 0.89 ± 0.22 | 2.91 ± 1.48 | 1.72 ± 0.21 | 1.54 ± 0.27 |
| | Ib H30656 | 1.43 ± 0.23 | 1.21 ± 0.16 | 1.64 ± 0.19 | 1.01 ± 0.21 |
| | FLR | 1.24 ± 0.09 | 1.86 ± 0.19 | 1.29 ± 0.18 | 1.13 ± 0.11 |
| | PRVABC59 | 0.69 ± 0.11 | 1.85 ± 0.51 | 1.25 ± 0.19 | 0.91 ± 0.2 |
| DENV | Type 4 H241 | 0.18 ± 0.12 | >50 | >50 | >50 |
| YFV | 17D | 0.27 ± 0.12 | >50 | >50 | >50 |
| JEV | SA-14 | 1.85 ± 1.1 | >50 | >50 | >50 |
| WNV | NY-99 | 3.00 ± 2.50 | >50 | >50 | >50 |

EC<sub>50</sub> values were measured in CPE-protection based assays with DENV type4 strain H241, JEV strains SA-14, WNV NY-99, and YFV strain 17D as described in the Methods and Materials. Data represent mean ± s.d. ( μM) from > 3 independent dose response analyses.

### 3. The MWAC-3475 series possesses excellent anti-ZIKV efficacy

Next, we used MWAC-3475 as a model compound for the characterization of the series in detail. Treatment of ZIKV-infected cells with MWAC-3475 resulted in significant reduction in progeny virus titers (**Figure 1**), >1000-fold reduction at 5 µM, which is comparable to that of NITD-008 (**Figure 1B**). The antiviral effect increased as the treatment concentration was increased, reaching a 4.2-log reduction (or 1.78 x 10^4^-fold) at 20 µM. The antiviral activity was further confirmed at the viral protein expression level by detecting viral E protein using the microscopy approach. The intensity and abundance of viral E protein detected with the 4G2 antibody decreased in a concentration-dependent manner. (**Figure 1C**).

**Fig 1.**
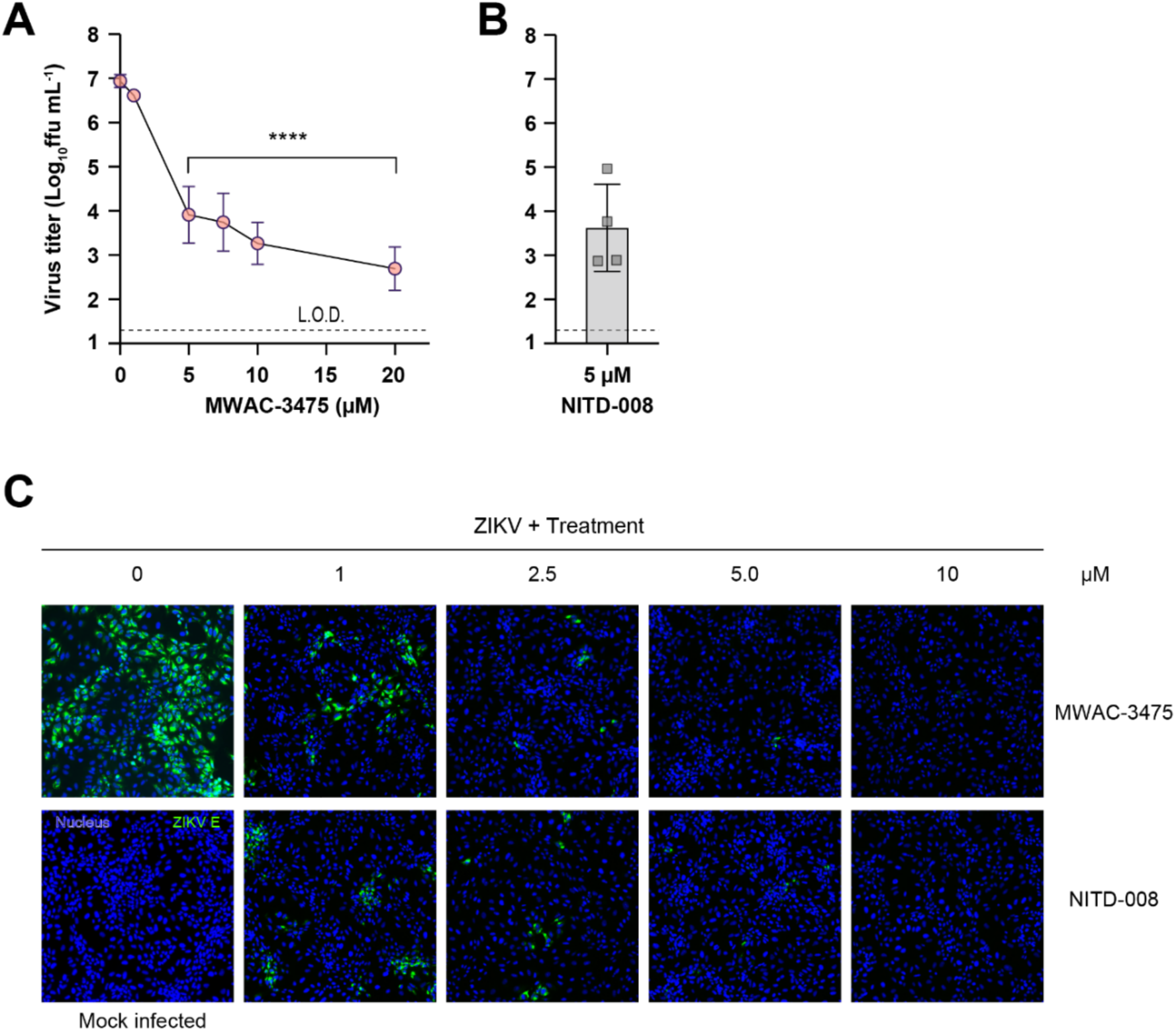
Anti-ZIKV activity of MWAC-3475. Vero76 cells were infected with ZIKV (strain PLCal, 0.05 m.o.i) and treated with various concentrations of MWAC-3475 or NITD-008. (A) and (B) titers of progeny viruses in the supernatants (n=4). **** P< 0.0001, One-way ANOVA Dunnett’s multiple comparisons test. (C) Decrease of the viral protein expression by MWAC-3475 treatment was evaluated with the indirect immunofluorescence assay with anti-E (clone 4G2) at DPI 2. Nucleus stain (blue) and anti-4G2 signal (green) were imaged using a fluorescence microscope.

### 4. Improvement of antiviral potency by medicinal chemistry

The potency, efficacy, ease of synthesis (from the addition of commercial *p*-(t-butyl) benzoyl chloride to commercial amino cyclopentane in the presence of a base), and small size of MWAC-3475 (MW 275) together justified further study and optimization efforts. We thus studied both commercial and internally prepared analogs to gauge structural features important for anti-ZIKV activity with low cellular cytotoxicity. Increasing size of the amide group from cyclopentyl to cyclohexyl to cycloheptyl gave increased anti-ZIKV activity (**Figure 2**), though MWAC-4001 showed appreciable cellular cytotoxicity. A commercial bicycloheptyl analog (MWAC-3508, tested as a mixture of four isomers) showed promise. Preparation of the separate endo- and exo-racemic compounds showed specificity for the exo-isomer (MWAC-3989). Several attempts to increase water solubility in the series by introducing heteroatoms into the amide region eroded antiviral activity (bottom row, **Figure 2**).

**Fig 2.**
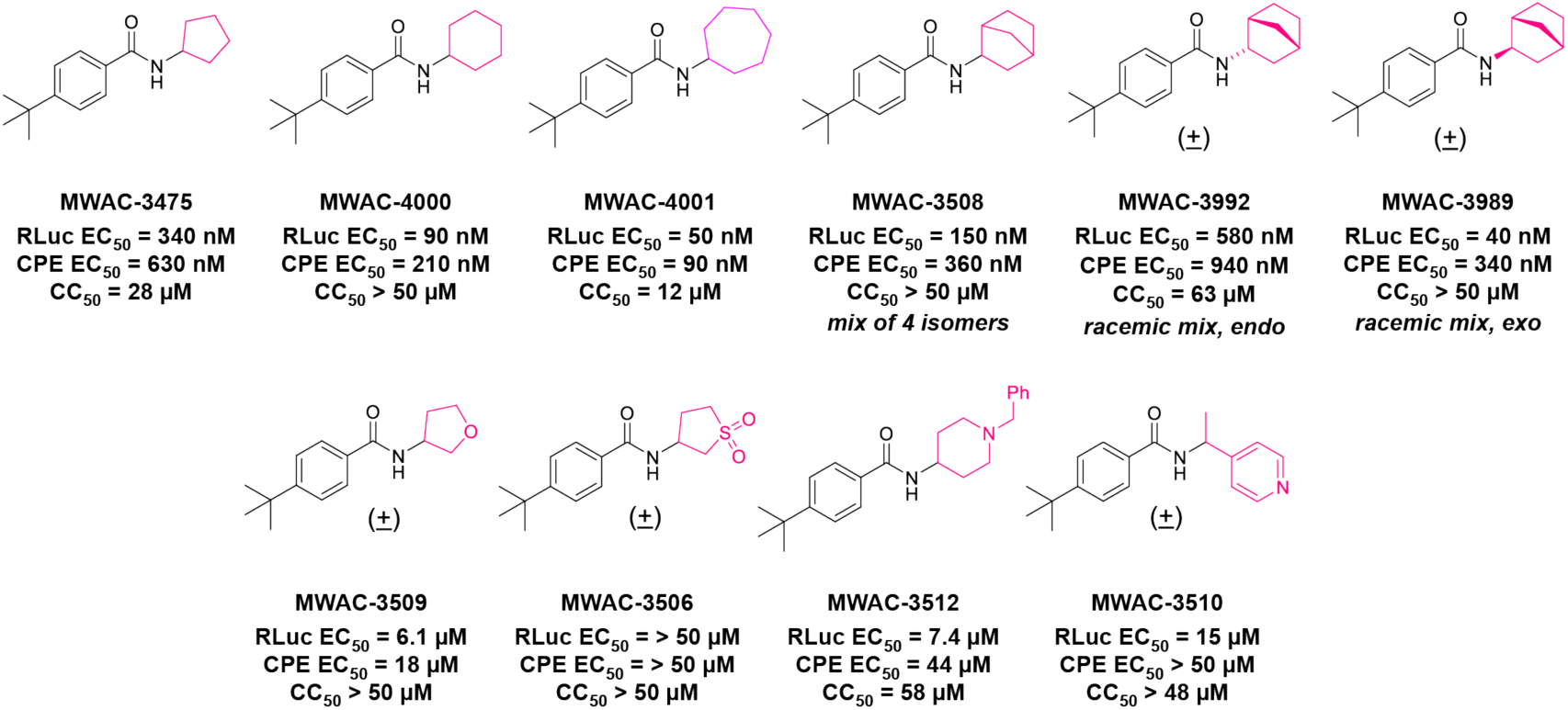
Amide region SAR studies.

Encouraged by the antiviral activity and low cytotoxicity seen for MWAC-3989, we retained this exo bicyclic amide group and probed the preference for the *para* substituent on the phenyl ring. In a focused set of seven analogs (**Table 3)** we found no suitable replacement for the *para*-t-butyl group. Further studies will broaden exploration in each region.

**Table 3.**
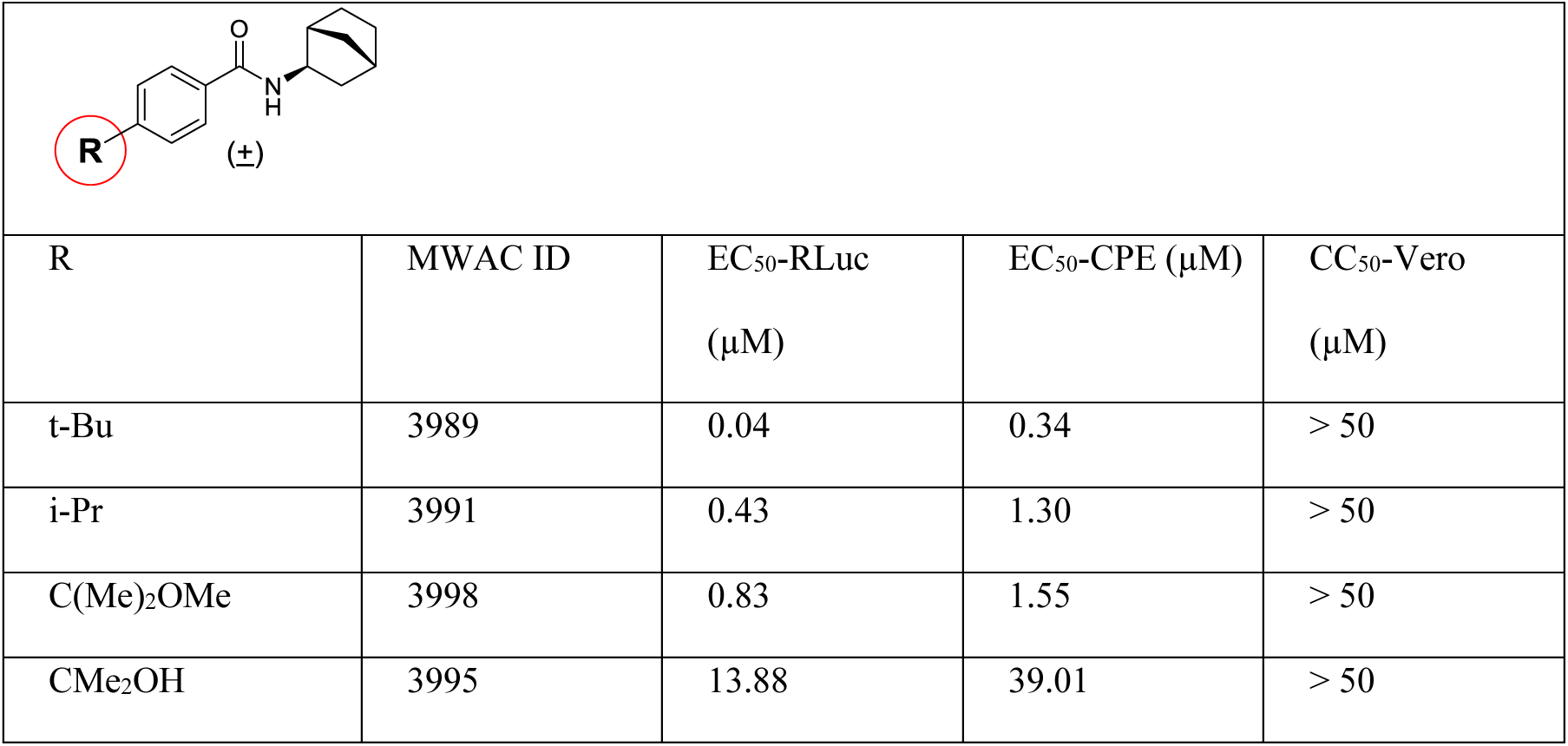
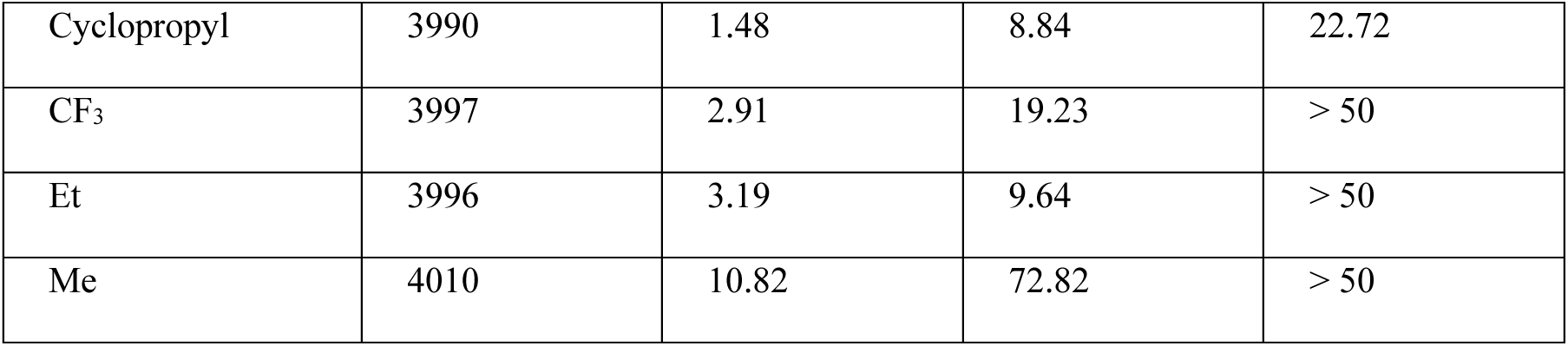
Modification of the phenyl substituent.

|  |  |  |  |  |
| --- | --- | --- | --- | --- |
| R | MWAC ID | EC<sub>50</sub>-RLuc<br>(μM) | EC<sub>50</sub>-CPE (μM) | CC<sub>50</sub>-Vero<br>(μM) |
| t-Bu | 3989 | 0.04 | 0.34 | > 50 |
| i-Pr | 3991 | 0.43 | 1.30 | > 50 |
| C(Me)<sub>2</sub>OMe | 3998 | 0.83 | 1.55 | > 50 |
| CMe<sub>2</sub>OH | 3995 | 13.88 | 39.01 | > 50 |
| Cyclopropyl | 3990 | 1.48 | 8.84 | 22.72 |
| CF<sub>3</sub> | 3997 | 2.91 | 19.23 | > 50 |
| Et | 3996 | 3.19 | 9.64 | > 50 |
| Me | 4010 | 10.82 | 72.82 | > 50 |

### 5. Preliminary in vitro DMPK assessment in the series

All analogs in the series were compliant with common druglikeness rules (e.g., Lipiniski’s and Veber’s rules, and their extensions)(28–30), but further *in vitro* assessments were warranted to validate the potential of the series for advancement. Properties and in vitro DMPK of the early bicycloheptane lead MWAC-3508 were analyzed (**Table S1**). Areas for improvement may include decreasing plasma protein binding, improving stability to liver metabolism in mice, and augmenting water solubility. Nevertheless, this lead and the series appear to have potential for further development.

### 6. Compounds in the lead series inhibit viral replication in the middle stage and require *de novo* protein synthesis

We conducted a time-of-addition study to determine the viral replication stage at which compounds in MWAC-3475 series intervene. We first measured viral RNA level in cells at various times after a synchronous infection with a high MOI (MOI=3) of ZIKV (**Figure 3A**). A significant accumulation of viral RNA synthesis initiated at 12 h.p.i. and peaked at 36 h.p.i, indicating the most productive phase of viral replication (i.e., the middle phase) occurs during this period. Detection of viral protein by western blot also confirmed the active viral protein synthesis during this period. (**Fig. S3**). When a test compound was added at different time point in relation to the infection, we found that its addition to infected cultures could be delayed, as long as active intracellular viral RNA synthesis had not initiated (i.e., at 12 h.p.i., **Figure 3B**) without loss of antiviral efficacy. When the inhibitor was added after onset of viral RNA synthesis (i.e., 16 h.p.i.), a gradual loss of its antiviral activity was noted, suggesting an interference with the viral RNA replication step in the middle phase of replication, which was similar to the pattern of NITD-008, a nucleoside analog vRNA synthesis inhibitor (**Figure 3B**). To understand if newly synthesized viral proteins are required for MWAC-3475’s antiviral action, we tested the effect of cycloheximide co-administration. Pre-treatment with cycloheximide in the middle of viral RNA synthesis abolished the antiviral effect of MWAC-3475, indicating the antiviral mechanism of MWAC-3475 requires new protein synthesis. These results indicate that MWAC-3475 series inhibits viral replication during the middle phase and that, presumably, *de novo* viral protein synthesis is required for its antiviral action.

**Fig 3.**
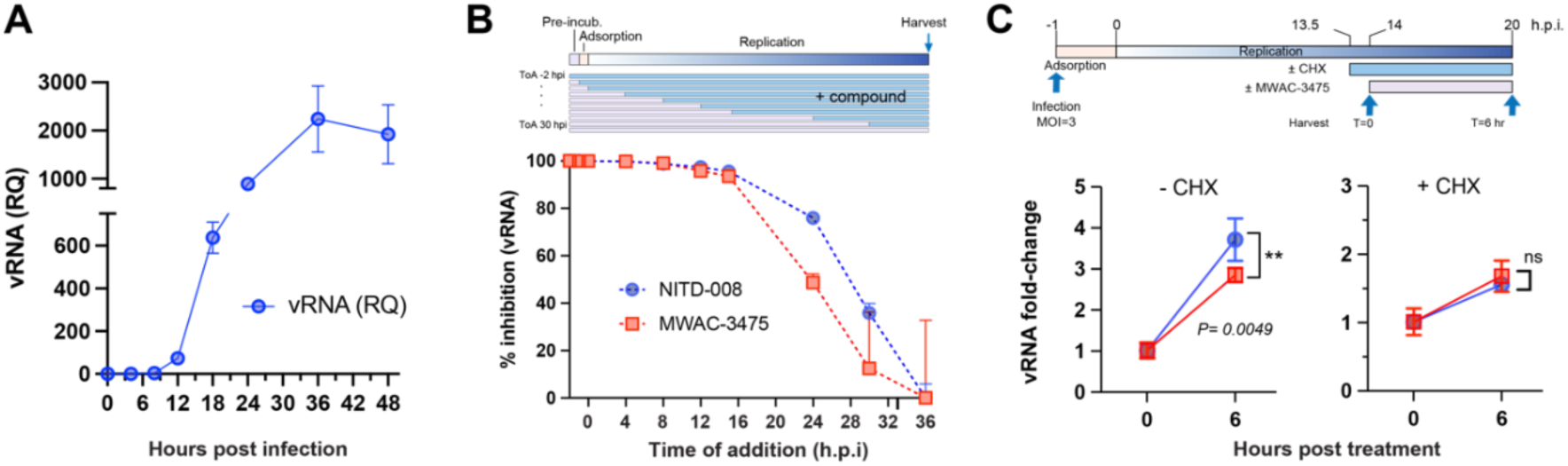
(A) Kinetics of viral RNA synthesis in ZIKV-infected cells. Vero cells infected with ZIKV (m.o.i. = 3) were harvested at times indicated (x-axis) and the viral RNA amount was quantitated with real-time PCR. (B). Time-of-addition. Compound was added at different point of times in respect to virus infection and harvested at 36 h.p.i. Viral RNA was measure using the real-time PCR and normalized using the 2^-ddCq^ method. (C). ZIKV-infected cells were treated with cycloheximide (CHX) 30 min. prior to the treatment of MWAC-3475 and incubated for 6 hours. Viral RNA was quantitated as described before.

### 7. Resistance studies implicate the viral protein NS4B as the molecular target of the MWAC-3475 series

Identifying mutants arising in selection of resistance studies is a useful method for target identification. Accordingly, we sought to select MWAC-3475-resistant isolates and to identify mutations conferring resistance. ZIKV was passed 10 times in the presence of MWAC-3475, gradually increasing concentration from 1 µM to 20 µM. During the serial passages the population maintained its infectivity, as high as > 10^7^ pfu/mL (**Figure 4A**). After the final passage the viral population showed resistance to MWAC-3475, as evidenced by the observation of plaque generation in the presence of the compound in the overlay media (**Figure 4B**). Sequencing of the passage 9 virus population identified MWAC-3475-specific mutations mapped within the C-terminal region of NS4B, aa 241-248 (V241L, A245T, and V248G. **Figure 4C**). These mutations map to the transmembrane domain 5 (TM-5). Interestingly, this site is distal from the site conferring resistance in DENV to previously developed DENV NS4B inhibitors (**Figure 4D**)(16–19). Sequence homology analysis of the identified residues (**Figure 4E**) showed a low consensus sequence homology across members of the flavivirus family, in accord with the ZIKV-specific activity we have observed (**Table 2**).

**Fig 4.**
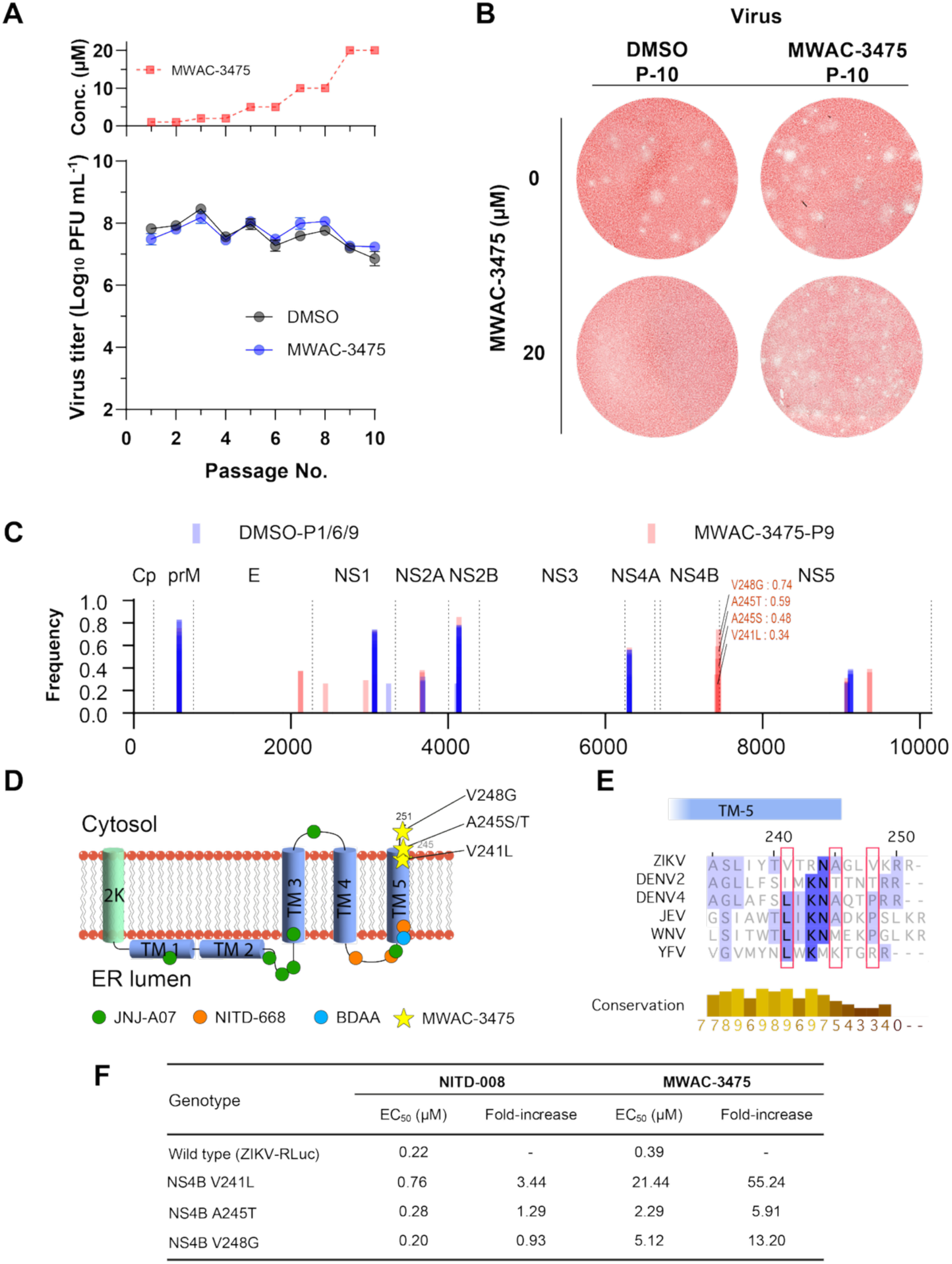
Identification of mutations conferring resistance to MWAC-3475. (A) Virus passage scheme with increasing concentrations of MWAC-3475 for 10 passages (top) and the viral titers at each passage. (B) Plaques of the 10th passage virus populations in DMSO or MWAC-3475 were developed in the presence (bottom) or the absence (top) of MWAC-3475 in the agarose-overlay media. After three days, cells were fixed and stained with neutral red dye. (C) Locations and frequencies of mutations of virus populations. Bars represent mutations identified from population passed in the presence of mock (DMSO, blue) and MWAC-3475 (red). Mutations of interest clustered within NS4B were noted with amino acid sequences and their frequency at passage 9. (D) locations of mutations conferring resistance to various anti-flavivirus inhibitors. 2K and transmembrane (TM) domains depicted as cylinders. Anti-DENV inhibitors (JNJ-A07 and NITD-668, green and orange circle), anti-YFV inhibitor (BDAA, cyan circle), and MWAC-3475 (yellow star) resistant mutations are depicted in the respect to their relative locations to each NS4B topology. (E) Sequence alignment around the C-terminal region of flavivirus NS4B. Red boxes represent the locations of amino acids related to the activity of MWAC-3475. (F) Phenotypic resistance by the NS4B mutations

For further validation we used a reverse-genetics approach, introducing these mutations into a molecular clone derived strain (ZIKV-RLuc) and measuring susceptibility to MWAC-3475. We found that introducing these mutations within NS4B resulted in a 6-to 55-fold decrease in potency of MWAC-3475 (**Figure 4F**). This phenotypic resistance by the mutations clearly indicates direct involvement of these residues of ZIKV NS4B in the antiviral activity of the series.

### 8. FAM fluorescence labeling provides further support for ZIKV NS4B as the target

While the phenotypic resistance study strongly implicates NS4B as the molecular target of the series, we sought further support using an *in situ* microscopy approach. We utilized a miniRFPnano3-tagged ZIKV NS4B and prepared a fluorescence-labelled analog of MWAC-3489, the racemic a-methyl *p*-methoxy benzyl derivative (**Table 1**). First, we prepared individual isomers and found that activity resides in the S-isomer MWAC-4168 instead of the R-isomer MWAC-4169, that the p-methoxy group is preferred over meta (MWAC-4170) or ortho (MWAC-4138) substitution, and that extension of the OMe group to a longer alkyl chain bearing a clickable alkyne handle (MWAC-3986) slightly augments rather than erodes activity (**Table S2**).

The anti-ZIKV activity and lack of cytotoxicity seen with MWAC-3986 justified a strategy of using the alkyne to link a FAM fluorophore to give a derivative suitable for direct imaging (see probe MWAC-4163, **Figure 5A**). 24 hours after delivery of the plasmid expressing the 2K-miniRFPnano3-NS4B polypeptide, cells were fixed, permeabilized, and incubated with MWAC-4163. The FAM fluorescence signal from the compound was shown in a punctuated pattern in cells without ZIKV NS4B expression (evidenced by miniRFPnano3). In cells with ZIKV NS4B expression, however, a strong colocalization between ZIKV NS4B (shown in red) and MWAC-4163 (shown in green) was detected specifically within the perinuclear region(31), indicating a strong specific interaction between them.

**Fig 5.**
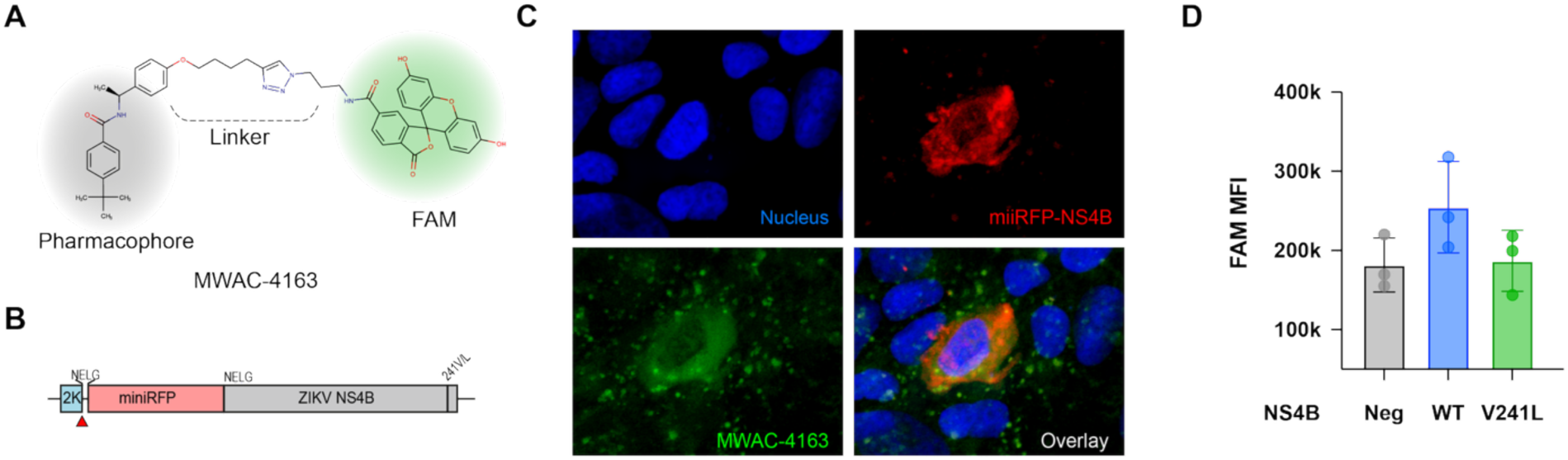
*In situ* interaction between the benzamide series and ZIKV NS4B. (A) Structure of MWAC-4163. (B-D). Colocalization of MWAC-4163 and NS4B. Plasmid encoding miniRFP-NS4B (B) was introduced into Vero cells for expression for 30 hrs. After fixation and permeabilization, cells were incubated with MWAC-4163 (10 µM), then imaged (C) with a confocal microscope. (D). FACS analysis showed a higher presence of MWAC-4163 in NS4B+ cells than in NS4B-cells. This phenotype was reversed by the V241L mutation.

To further investigate if the interaction could be mediated by the C-terminal region of the NS4B, as indicated from the resistant mutant study (**Figure 4**), we conducted a similar experiment with the same construct with the mutations (e.g., NS4B V241L or V248G) conferring resistance to the series. Cells expressing NS4B with these mutations showed a lower degree of co-staining than that of the wild type NS4B. This finding confirmed that the amino acid residues in the C-terminal region of the NS4B are responsible for the interaction with the lead small molecule series.

### 9. ^19^F NMR studies provide further support for ZIKV NS4B as the target

The flavivirus protein NS4B is membrane bound(32), a property that has hindered structure elucidation for complexes of DENV NS4B(33) with clinical anti-DENV compounds found to target NS4B. ^19^F-NMR spectroscopy can be used with such targets, however, and was the method we chose for studying interactions of fluorine-containing small molecules with ZIKV NS4B. We first validated the method using DENV NS4B and JNJ-1802, a fluorine-containing pan-DENV NS4B inhibitor(34). When the ^19^F-NMR spectra of JNJ-1802 (at 50 µM) combined with NS4B (5 µM) was compared to that of free JNJ-1802, a chemical shift perturbation and significant line broadening was seen, which are both strong evidence for direct interaction (**Figure 6A**). A similar study using our series of anti-ZIKV compounds required a fluorinated analog, MWAC-3533, which has modest anti-ZIKV activity (RLuc EC_50_ ∼16 μM). ^19^F-NMR signals for this compound were shifted by the presence of ZIKV NS4B in a concentration-dependent manner (**Figure 6B**). Ionic strength/pH was not significantly altered in dilutions (data not shown). K_d_ determination using ^19^F shifts in general aligned with the activity seen in the RLuc assay (**Figure 6C**). Together these observations further support the hypothesis that this series directly binds to and inhibits the function of ZIKV NS4B.

**Fig 6.**
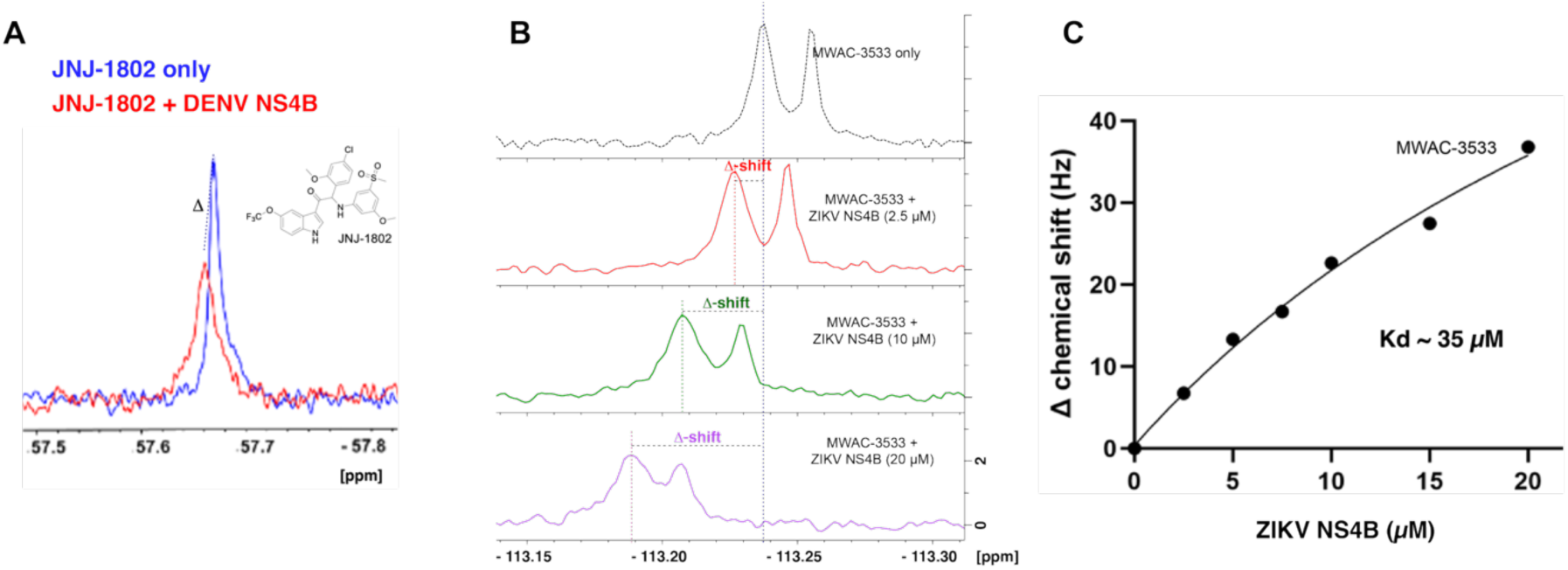
^19^F-NMR studies. (A) Overlaid ^19^F-NMR spectra of JNJ-1802 in absence (blue) and presence of DENV NS4B (red). Both chemical shift perturbation and line broadening are evidence for target engagement. (B). Interaction between ZIKV NS4B and MWAC-3533 shown in ^19^F-NMR. (C). Measurement of K_d_ through titration and chemical shift perturbation analysis.

### 10. Investigation of drug synergy potential: NS4B inhibition by MWAC-3475 may synergize with viral polymerase inhibition

Combinations of antiviral drugs acting on different but critically important pathways are attractive with respect to robust and sustained efficacy with reduced resistance potential. Accordingly, we surmised that targeting ZIKV NS4B in combination with another anti-ZIKV agent with an unrelated mechanism of action would be an effective strategy. To quantify multi-drug effects we used the concentration-responsive, CPE-based anti-ZIKV assay and evaluated synergy scores for a combination of MWAC-3475 with NITD-008, a nucleoside analog targeting the viral polymerase (**Figure 7**). Overall, the mean synergy scores were relatively moderate (e.g., ZIP score 5.47); however, we found a significant synergistic effect between the two compounds within a concentration range near their EC_50_ for both compounds, with ZIP and HSA synergy scores of 42.08 and 48.08, respectively (**Table S3)**. Combination significance scores were high (>79), indicating a potential synergistic or at least a robust additive effect from the combination(35,36). When MWAC-3475 was instead tested together with an analog in the same series, synergy (as expected) was not observed (not shown).

**Fig 7.**
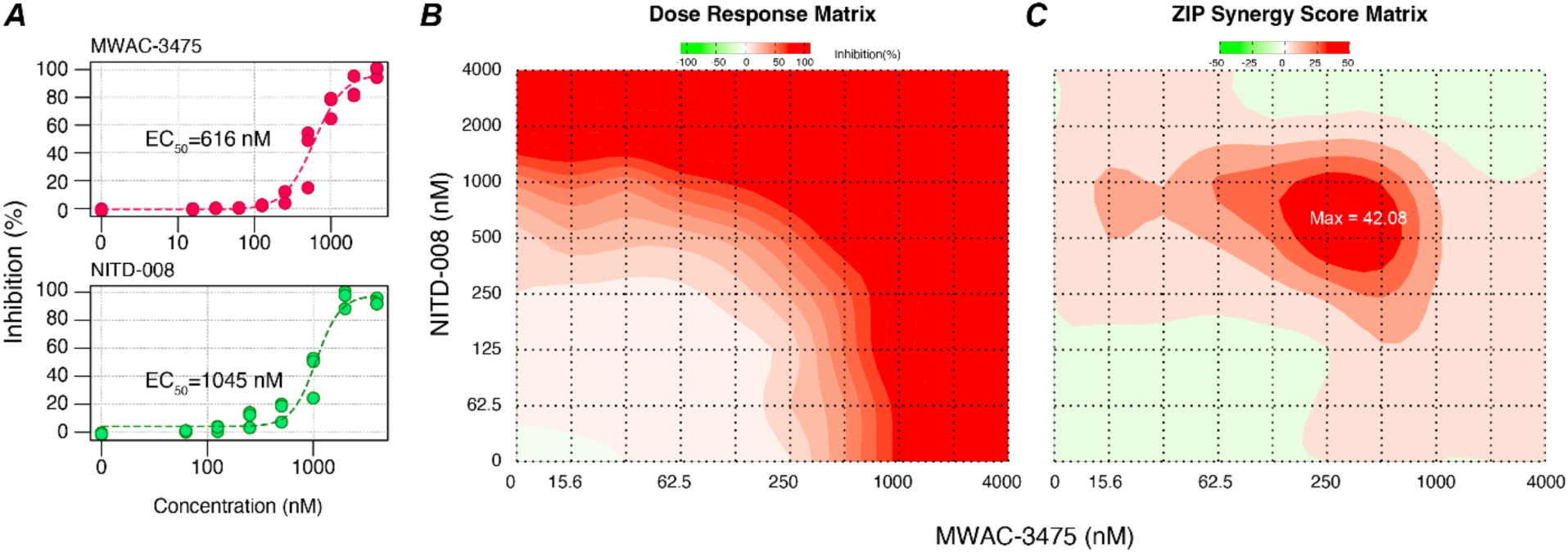
Synergistic effects for a combination of MWAC-3475 with NITD-008. (A) Performance of MWAC-3475 and NITD-008 as a single agent in the dose-responses anti-ZIKV assay. (B) Dose response of antiviral effect at various concentration matrix (n=3, per point). (C) Contour diagram of the predicted ZIP synergy scores. The analysis and graphs were generated by using SynergyFinder Plus without a baseline correction.

## 5. DISCUSSION

Since the “re-emergence” of ZIKV around 2014 in Asia and the Americas, ZIKV has become endemic at a global scale. In Americas between 2022 and 2025, an average of approximately 40,000 cases of ZIKV have been reported per year, with highest case numbers in Brazil and neighboring countries. ZIKV infection is particularly concerning due to its devastating outcomes to infants born to mothers infected by ZIKV during the pregnancy. Approximately 5% of infants born to mothers infected with ZIKV during pregnancy develop Zika-associated birth defects, including microcephaly and other various developmental deficits(37). The horizontal transfer of ZIKV from mother’s blood to fetal brain has been shown to drive these complications(38,39). Hence, an effective treatment to substantially lower ZIKV levels in the mother would be expected to avert development of congenital Zika syndrome in the baby, a premise supported by results from a monoclonal antibody treatment in mouse models(40). In addition, infected humans are a critical contributor to ZIKV urban transmission. Therapeutic or preemptive prophylactic treatments of people in epidemic regions could lower the risk of viral transmission to pregnant mothers and further contribute to curtailing of viral spread. However, no effective therapeutics or potential candidates are currently available for ZIKV. Further studies are warranted to ascertain the most effective treatment strategies, such as acute vs. prophylactic treatments. With further study, compounds related to the agents here disclosed may show promise, alone or in combination with other antivirals, in thwarting the spread of ZIKV outbreaks.

In this study we developed a phenotypic HTS assay to identify antiviral candidates, leading to the discovery of benzamide-based hit compounds. CPE-based antiviral HTS efforts can effectively identify novel antiviral compounds with a low cytotoxicity. However, one of the drawbacks is selecting compounds that interfere with cellular (host) targets, which may be liable for potential toxicity as a hit. Particularly, cellular pyrimidine synthesis inhibitors have been reported often from antiviral HTS campaigns by our group and others (14,24–26,41–43). We circumvented the problem by supplementing the assay media with uridine, a precursor for the pyrimidine salvage pathway, which could neutralize the inhibitory effect of a compound through the de novo pyrimidine synthesis pathway (e.g., DHODH). Our strategy seemed to have worked as evidenced by the fact that only one compound from the 91 compounds selected for the concentration-response assay (Figure 1) showed a non-ZIKV specific antiviral activity (EC_50_-CHIKV = 6 µM).

The initial hits demonstrated selective activity against ZIKV, through across ZIKV strains. Subsequent optimization yielded potent derivatives with favorable physical and in vitro DMPK properties, supporting their potential as antiviral agents. To elucidate the molecular target of this compound series, several viral isolates resistant to MWAC-3475 were characterized, giving evidence that NS4B is its target. Experiments with viruses harboring the identified NS4B mutations also showed phenotypic resistance, further confirming NS4B as the target responsible for the antiviral activity of our series. To probe direct interactions between the compounds and NS4B, we studied a fluorinated analog by ^19^F NMR spectroscopy, using recombinant NS4B in detergent micelles, a method first validated with DENV NS4B and JNJ-1802. To reduce the influence from the micelles, ^19^F-NMR was used in probing protein-inhibitor interactions as fluorine atoms are not present in biological systems. Clear evidence for a direct interaction between ZIKV NS4B and the compound was observed, further confirming NS4B as the target. Taken together, our study demonstrates that a robust workflow including phenotypic screening, compound optimization, target identification through generating resistant viruses, and NMR structural biology can lead to antiviral compounds for development.

Previously NS4B has been reported as the antiviral target of hits discovered by other phenotypic assay-based HTS campaigns for flaviviruses, such as JNJ-A07(34,44), NITD-618(45), Compound 14a(46), NITD-668(19) for DENV, and BDAA series(18) for yellow fever virus. It is quite intriguing to see NS4B as a convergent target from these successful various flavivirus antiviral discovery campaigns, despite large differences in chemical structures and the phenotypic resistant profiles of each. This could be due to the multifaceted biological roles of NS4B. Flavivirus NS4B is a membrane protein without known catalytic activity. It forms homomultimers(47–49) and interacts various viral non-structural proteins, including NS1(50), NS2B(51), NS3(52–54), NS4A(55), and host proteins(56) (see a review article for the comprehensive list of proteins(57)). NS4B is expected to play critical roles in formation of viral replication organelles (ROs) and to serve as a foundation for building ROs for RNA synthesis. These diverse interactions with critical viral proteins centered around NS4B could provide multiple druggable sites, leading to identify multiple hits targeting various regions of NS4B. These NS4B inhibitors including our series shares common aspects such as 1) high specificity to the target virus, and 2) inhibition of *de novo* formation of the viral replicase complex(44) or membrane remodeling(58), resulting in effective viral RNA synthesis inhibition. However, the resistant mutations are specific to the corresponding viruses, indicating the mode of action at the molecular level might be unique for each antiviral and its target NS4B.

The benzamide series-resistant mutations (a.a. positions of 241, 245, and 248) were found between the terminus of the potential transmembrane domain 5 (pTMD5) and the C-terminus of NS4B(a.a. 251)(49,59). Interestingly this region in DENV NS4B has been reported important for stable expression of NS4B on the ER via interaction with the ER membrane complex (EMC)(60). DENV2 with NS4B N246Y (i.e., A/T mutation at 7558 n.t.) mutation was able to restore the replication ability in EMC4-deficient cells, implying the region might be critical to interact with the EMC for proper topology of potential transmembrane domains (pTMDs). A follow up mechanism of action study will be warranted to understand how the series interacts with ZIKV NS4B, and how the interactions result in the inhibition of viral replication in detail.

NS4B, a critical protein for replication of flaviviruses (an emerging family of RNA viruses of high pandemic potential), had been validated as an antiviral target with clinical anti-DENV compounds. Thus, it is not surprising that perturbation of ZIKV NS4B function inhibits ZIKV replication, though no prior reports of ZIKV NS4B inhibitors have emerged. Given that NS4B is involved in various functions, including protein-protein interactions, we also explored combination strategies to enhance antiviral efficacy. MWAC-3475 showed synergistic effects when used with a viral polymerase inhibitor but not with structurally similar compounds. This suggests the potential of combining NS4B inhibitors with agents targeting different pathways, focusing on enzyme inhibition of proteins such as NS3 and NS5, which often exhibit low druggability. Our discovery once again demonstrated NS4B as a vulnerable point (i.e., Achilles heels) for flaviviruses, establishing it as the valid antiviral target. Despite being an integral membrane protein lacking enzymatic activity, which complicates structure-based drug discovery, our findings provide a framework for overcoming these challenges. A well-designed screening strategy is critical for identifying active compounds and confirming the activity of the developed compounds. Iterative medicinal chemistry optimization is indispensable for improving potency and pharmacological properties. Determining the molecular target is crucial for understanding the mechanism of action and for designing potential combination treatment strategies. Both biochemical and biophysical validation assays are necessary to confirm direct compound-target interactions. Despite the process made in current studies, further structural characterization of NS4B– inhibitor complexes will provide binding mechanism of this series of inhibitors and support rational design of next-generation antivirals.

We herein present a novel small molecule series that inhibits ZIKV replication by interfering with the NS4B. Importantly, our study establishes NS4B as a valid antiviral target for flaviviruses. However, so far none of the reported NS4B inhibitors showed a pan-flavivirus activity, indicating a challenge in developing a broad-spectrum antiviral for flavivirus targeting the NS4B. A future study to develop the benzamide series into a therapeutic candidate for ZIKV as well as for pan-flaviviruses would be warranted.

## 6. MATERIALS AND METHODS

### Virus and Cells

Vero 76 Cells (ATCC® CRL-1587™) were maintained in Modified Eagle’s Medium with Earle’s Balanced Salt Solution and L-glutamine (MEM-E) supplemented with 10% fetal bovine serum (FBS) (Corning CellGro). Cells were maintained at 37 ℃ in humidified incubators with 5% CO_2_. Virus stocks were obtained from BEI resources and amplified once before being used for experiments. Amplified stock virus was aliquoted and stored at -80 °C until used.

The ZIKV strains MR766 and PRVABC59 (GenBank: KX087101) were provided by Dr. Barbara Johnson at the CDC. The following reagents were obtained through BEI Resources, NIAID, NIH: ZIKV strains of PLCal_ZV (Human/2013/Thailand; GenBank: KF993678; BEI NR-50234), Ib H30656 (GenBank:HQ234500: BEI NR-50066), FLR (GenBank: KX087102; BEI NR-50183), DENV Type4 (H241/TC; BEI NR-86), and YFV strain 17D (BEI NR-116). DAK-41525-MA was a gift from Dr. Nair. CHIKV 181/25 was rescued from the pCHIKV181/25 plasmid (a gift from Dr. Sokoloski) following a procedure described previously (PMID: 39448734) A plasmid pZIKV-RLuc was a gift from the World Reference Center for Emerging Viruses and Arboviruses, The University of Texas Medical Branch. It has the cDNA of Zika virus isolate FSS13025 genome (GenBank: MN755621.1) and the Renilla luciferase gene as a reporter. The whole genome and reporter cassette was subcloned into the CopyControl pCCBAC vector by using a standard PCR-based cloning approach. Plasmid (pCC-ZIKV-RLuc) linearized with FesI was used as a template for in vitro RNA synthesis using (mMessage mRNA synthesis) following the manufacturer’s protocol with addition of 2 µL of GTP mix in a 20 µL reaction. Produced RNA was purified and approximately 5 µg of RNA was transfected to Vero 76 cells grown in a 6-well plate with Lipofectamine MessengerMax (Thermo Fisher Scientific Inc.). Rescued virus (ZIKV-RLuc hereafter) was harvested 5 days post transfection and was amplified once for use. Virus titration was done by the TCID-50 method based on the RLuc expression with the Renilla-Glo assay system (Promega, USA).

Specific mutant viruses were generated using the Quick-change mutagenesis with the pCC-ZIKV-RLuc template (See Supportive information for the primer sequences). The full sequence of the resultant clone was validated prior to rescuing the virus. Virus rescue was performed as described above.

### Antiviral assays

Compounds were dissolved in DMSO at 20 mM and stored at -20 °C. Antiviral potency assays (i.e., EC_50_ assays) were done in the 96-well format. Briefly, Vero 76 cells seeded in a 96-well plate one day prior were infected with virus at a virus-specific optimal MOI in the presence of test compounds, which were serially diluted to 8 different concentrations. The final concentration of DMSO was maintained at 0.25%. For CPE-based assays, infected cells were incubated for virus-specific optimal incubation times prior to measuring cell viability with CellTiter-Glo (Promega) to assess protection from virus-induced CPE. A different combination of an MOI and an incubation time that was optimized for consistent and robust performance was used for each virus assay. For ZIKV, an MOI of 0.05 and a 3-day incubation was used while for YFV 17D and DENV, an MOI of 0.05 and 0.1 was used with an incubation time of 4 and 6 days, respectively.

### CPE-based phenotypic high-throughput screen

To begin, 600 cells at 2.5 μL / well of Vero 76 cells in MEM-E supplemented with 10% FBS and 1X Antibiotic-Antimycotic (Gibco 15240-062) were dispensed to 1536 well plate (Aurora EBB0-40000A) and incubated at 37 ℃ in humidified incubators with 5% CO_2_ for 24 hours. After that, 30nL of each compound or vehicle (75% DMSO) were transferred using pintool and further incubated for 2 hours. Then, 2.5 μL / well of ZIKV PLCal at MOI 0.5 in virus dilution medium (MEM-E supplemented with 10% FBS, 12.5 mM HEPES, 100 μM uridine and 1X Antibiotic-Antimycotic) was dispensed. For the negative control wells (N=24/plate), only virus dilution media was dispensed. Positive control wells contained virus and cells (N=24/plate). After incubation at 37 ℃ in humidified incubators with 5% CO2 for 72 hours, plates were incubated for 10 minutes at room temperature, followed by dispensing 5 μL/well CellTiter Glo (Promega). Luminescent signal was detected using PHERAstar FSX (BMG Labtech).

In the primary screen, compounds were tested at a single concentration in singlicate at a final nominal concentration of 6.95 μM. Raw assay data was imported into UF Scripps’ corporate database and subsequently analyzed using Symyx software. Activity of each compound was calculated on a per-plate basis using the following equation: Percent Response of compound= 100 * ((Test Well-Median Data Wells) / (Median High Control - Median Data Wells))

A summary of the results of the primary screening assay is shown in **Fig. S2**. A mathematical algorithm was used to determine active compounds on a per plate basis. Three values were calculated: (1) the average activity value for all sample wells (2) 3 times the stdev value for the same set of wells (3) The sum of these two values was used as a cutoff parameter, i.e. any compound that exhibited greater percent inhibition than the cutoff parameter was declared active; again, this was applied to each individual plate. For the confirmation assay we applied the same formula to determine the hit cut-off. For the titration assays, a four-parameter equation describing a sigmoidal dose-response curve was then fitted with adjustable baseline using Assay Explorer software (Symyx Technologies Inc.). The reported EC_50_ values were generated from fitted curves by solving for the X-intercept value at the 50% inhibition level of the Y-intercept value. Compounds with an EC_50_ greater than 10 μM were considered inactive, while compounds with an EC_50_ less than 10 μM were considered active.

### Test compounds

The following compounds, listed in order of appearance in the manuscript, were purchased from the indicated vendor and tested as provided: MWAC-3400 (ChemDiv, K781-6490), MWAC-3415 (Chembridge, 7435251), MWAC-3417 (Chembridge, 7268948), MWAC-3481 (Life Chemicals, F1175-0153), MWAC-3489 (Vitas-M Labs, STL197485), MWAC-3475 (Enamine, Z32413960), MWAC-3508 (Chembridge, 6486655), MWAC-3509 (Chembridge, 7945795), MWAC-3506 (Chembridge, 5551926), MWAC-3512 (Chembridge, 9297289), MWAC-3510 (Chembridge, 9054690), and MWAC-3533 (Chembridge, 69155285).

Non-commercial compounds were prepared by direct amide coupling of a *para*-substituted benzoic acid, such as 4-t-butyl benzoic acid, with a commercially available amine, or alternatively by acylating the amine with a *para*-substituted benzoyl chloride (see Supplementary material).

### Property Assessments of MWAC-3508

Calculated properties (e.g., MW, LogP, tPSA) were assigned based upon input structure by the CDD (Collaborative Drug Discovery) Vault database using accepted algorithms. In vitro property assessments (CYP450 inhibition, kinetic solubility, liver microsomal stability, plasma protein binding) were conducted by WuXi AppTec, Inc. following established protocols.

### Western blot

ZIKV-infected Vero 76 cells were harvested at various points post infection and suspended in 200 µL of RIPA buffer (50 mM Tris-HCl, 150 mM NaCl, 1% Triton-X100, 0.1% SDS) for lysis. Proteins in the lysates were separated by SDS-PAGE and transferred on to PVDF membranes (BioRad). After a blocking with 5% non-fat dry milk and 1% of normal goat serum in TBS-T for one hour, blots were probed with following primary antibodies in TBS-T with 5% non-fat dry milk overnight: anti-ZIKV NS4B (Invitrogen MA5-47129 1: 500), anti-ZIKV NS5 (Invitrogen PA5-143441, 1: 500), anti-ZIKV NS3 (GeneTex, GTX13309, 1: 2000), anti-actin (Cell Signaling, 12262S, 1:1000). After washing in TBS-T, blots were incubated with HRPO-conjugated goat anti-rabbit secondary antibody (Santacruz, Sc-2054, 1: 10,000) then imaged with chemiluminescent substrate using an imager (Azure).

### Realtime PCR

Total RNA from the infected cells were isolated by using RNAzol-RT (MRC, Inc.) following the manufacturer’s instruction. RNA was suspended in THE RNA Storage Solution (Invitrogen) in 50 µL/sample, and RNA was reverse transcribed to cDNA with Maxima reverse transcriptase primed using a mixture of random hexamers (0.5 µM) and oligo-dT (0.31 µM) following the manufacturer’s protocol. cDNA was subjected to real-time PCR in a multiplex format with 2X TaqMan Gene Expression Master Mix (ABI), 500 nM of ZIKV-specific primers and 250 nM of probes (FAM-tagged) (61), and a Vero GAPDH-specific primer/probe set (SUN-tagged, See Supplementary information for sequences). Realtime PCR was performed with Quanta7Pro (ABI) and analyzed by using the 2^-ddCq^ method with three biological replicates and three technical replicates.

### Time-of-Addition study

Vero 76 cells cultured in 6-well plates overnight were infected with virus at an MOI of 3 for one hour. After removal of virus, cells were rinsed with PBS and replenished with fresh media (T=0). Compound (10 µM of MWAC-3475) or DMSO was added at the times indicated in the study and the total cells were harvested at 36 h.p.i. for total RNA isolation. Quantitation of viral RNA was performed as described above with three biological replicates.

For cycloheximide treatment, cells were started being treated with cycloheximide (Sigma) at 12.5 µg/mL or DMSO (n=3) at 13.5 h.p.i. At 14 h.p.i., test compound was added, and the infected cells were incubated at 37 °C for 6 hours (i.e., 20 h.p.i.). Total RNA was harvested from the cells and subjected to real-time PCR for RNA quantitation as described above.

### Serial passage and mutant virus sequencing

For the initial infection, Vero 76 cells grown in 6-well plates were infected by incubating the cells with ZIKV PLCal at MOI of 1 for one hour at 37 ℃. The cells were washed with PBS and replenished with virus growth media containing either MWAC-3475 or DMSO. 0.125mL of the cell supernatant was harvested to blindly inoculate for the next round of infection. The remainder of the passages followed the procedure described above. The progeny virus titers were enumerated as described above, and infected cells were resuspended in 1mL RNAzol RT (Molecular Research Center, Inc.) for total RNA purification following the manufacturer’s protocol. The cDNA was synthesized by using approximately 2 µg RNA whose secondary structures were removed by heating to 65°C for 5mins and cooling on ice for 5 more mins, 2µM random hexamers (IDT), RT Buffer, RNaseOUT Ribonuclease Inhibitor, 0.5mM dNTPs, and Maxima H-minus Reverse Transcriptase (Thermo Scientific). The ZIKV genome was PCR amplified by employing the ZIKV tiling strategy; with 0.5µM of different primer pools(62), 0.2mM dNTPs (IDT), and Q5 High-Fidelity DNA Polymerase and Buffer (NEBNext) on a S1000 Thermal Cycler (Bio-Rad) utilizing a protocol of: 98°C for 30s, 35 cycles of 98°C for 15s and 65°C for 5min. The amplicons from each sample were pooled and bead purified using KAPA Pure magnetic beads with a 0.8x bead-to-sample ratio (Roche) following the manufacturer’s protocol. The samples from each passage were combined to 1 mg for sequencing library preparation using the Nanopore Ligation Sequencing kit and Native Barcoding Expansion kit. Sequencing was performed using MinION Mk1b and Mk1c sequencers with R9.4.1 Flonges for each passage for 18 hr with high accuracy basecalling (Oxford Nanopore Technologies). The demultiplexed fastq files that that met the Phred quality score threshold were aligned to the reference genome sequence (ZIKV PLCal GenBank: KF993678.1) using Minimap2(63) {REF: PMID 29750242}, and mutations and their frequencies were identified by using mpileup from samtools (now bcftools) (64). For the sequenced samples, the average sequencing depth was 2632x, and the average coverage was 99.90%.

### Construction of NS4B expressing plasmids and colocalization analysis

First, pCDNA-miniRFP-NS4B was constructed by inserting the ZIKV NS4B gene at the C-terminus of the miniRFPnano3 gene in pCDNA-miniRFPnano3 plasmid (Addgene plasmid No. 184664)(65) by using the Gibson cloning method. Each part was PCR-amplified by using Phusion thermostable polymerase (NEB) with primers with overlapping sequences. Then chemically synthesized 2K gene (IDT DNA) was inserted into pCDNA-miniRFP-NS4B to construct pCDNA2K-miniRFPnano3-NS4B. The sequences of the final construct were verified by the whole plasmid sequencing. For expression, the plasmid was delivered to Vero 76 cells grown in cover slips by using Lipofectamine 3000 followed with the manufacturer’s protocol. At 24-hours later, cells were fixed with 2% paraformaldehyde and permeabilized with 0.1% Tween-20 in PBS for 20 min. Then, cells were incubated with MWAC-4163 at 20 µM in PBS for 30 min, followed by washing and nucleus counter staining with Hoest 33342. After mounting with a mounting media (ProLong Gold Antifade Mountant with DNA Stain DAPI, ThermoFisher), slides were imaged using a confocal microscope (Zeiss, LSM710 AxioObserver with a 63x objective).

### Purification of ZIKV NS4B

Purification of ZIKV NS4B was performed using the method described previously with slight modifications. The cDNA encoding full-length ZIKV NS4B was cloned into pET15b and transformed into *E. coli* BL21(DE3) Rosetta T1R cells. The cells were grown in 2xYT medium, and protein was induced by adding 0.5 mM β-D-1-thiogalactopyranoside (IPTG). The cells were then cultured at 18 °C, 180 rpm for 24 hours. Recombinant protein was then purified in the presence of lyso-myristoyl phosphatidylglycerol (LMPG) micelles as described previously(66). Protein was flash-frozen and stored in the SEC buffer containing 20 mM HEPES pH 8.0, 150 mM NaCl, 0.05% LMPG, 2 mM DTT at -80 °C.

### 19F-NMR experiments

To investigate the molecular interactions between NS4B and the test compounds, ^19^F-NMR experiments were performed on a Bruker spectrometer operating at a proton frequency of 400 or 700 MHz and equipped with a BBO probe and a cryoprobe. All measurements were conducted at 298K using standard pulse sequences from the Bruker pulse program library. ^19^F spectra were recorded for the compounds at a concentration of 50 µM. The changes of the spectra in the absence and presence of NS4B were recorded to probe the molecular interactions. To estimate the binding affinity, a titration experiment was conducted at a fixed concentration of 50 µM MWAC-3533 in NMR buffer (20 mM HEPES pH 8.0, 150 mM NaCl, 0.1% LMPG, 2 mM DTT, 5% DMSO, 10% D_2_O) in vary ZIKV NS4B concentration of 0, 2.5, 5, 7.5, 10, 15, and 20 µM. The resulting concentration-dependent chemical shift perturbations were analyzed using global nonlinear regression(67).

### Combination treatment and synergy scoring

Antiviral effect was tested in antiviral assays with combinations of an 8 × 10 dose matrix format, with one column and row dedicated for a single compound treatment. Each were serially diluted 2-fold by starting from a top concentration of 4 µM. CPE-protection activity was calculated as above and synergy effect was calculated by using SynergyFinder R package (version 3.16) without baseline correction(35,36).

## 8. ACKNOWLEDGMENTS

## Author contributions

Conceptualization : DC,LS,TPS,TDB

Data curation: DC,JS,LS,TS,TDB

Formal analysis: DC,TPS

Funding acquisition: DC,LS,TPS,TDB

Investigation: YO,EK,NMK,NKH,JK,RT,BA,KB,JM,JS,SU,CFT,YBT,CSC,CBK

Methodology: DC,YO,JK

Project administration and supervision: DC,LS,TPS,TDB

Validation: DC

Visualization: DC, TDB

Writing – original draft: DC,CBK, DL,TPS,TDB

Writing – review & editing: DC,DL,TPS,TDB

## Data and materials availability

All data are available in the main text or the supplementary materials.

## Supplementary Materials

Attached as a separate file.

## REFERENCES

1. Zardini A, Menegale F, Gobbi A, Manica M, Guzzetta G, d’Andrea V, et al. Estimating the potential risk of transmission of arboviruses in the Americas and Europe: a modelling study. The Lancet Planetary Health. 2024 Jan;8(1):e30–40.

2. Xu Y, Zhou J, Liu T, Liu P, Wu Y, Lai Z, et al. Assessing the risk of spread of Zika virus under current and future climate scenarios. Biosafety and Health. 2022 June;4(3):193–204.

3. PAHO. Cases of Zika virus disease [Internet]. Pan American Health Organization; Available from: https://www.paho.org/en/arbo-portal/zika-data-and-analysis/zika-analysis-country

4. Zika epidemiology update - May 2024 [Internet]. [cited 2025 June 25]. Available from: https://www.who.int/publications/m/item/zika-epidemiology-update-may-2024

5. Giraldo MI, Gonzalez-Orozco M, Rajsbaum R. Pathogenesis of Zika Virus Infection. Annu Rev Pathol. 2023 Jan 24;18:181–203.

6. Musso D, Ko AI, Baud D. Zika Virus Infection — After the Pandemic. Longo DL, editor. N Engl J Med. 2019 Oct 10;381(15):1444–57.

7. Rasmussen SA, Jamieson DJ, Honein MA, Petersen LR. Zika Virus and Birth Defects — Reviewing the Evidence for Causality. N Engl J Med. 2016 May 19;374(20):1981–7.

8. Krauer F, Riesen M, Reveiz L, Oladapo OT, Martínez-Vega R, Porgo TV, et al. Zika Virus Infection as a Cause of Congenital Brain Abnormalities and Guillain-Barré Syndrome: Systematic Review. PLoS Med. 2017 Jan;14(1):e1002203.

9. Vasilakis N, Weaver SC. Flavivirus transmission focusing on Zika. Current Opinion in Virology. 2017 Feb;22:30–5.

10. Barzon L, Pacenti M, Franchin E, Lavezzo E, Trevisan M, Sgarabotto D, et al. Infection dynamics in a traveller with persistent shedding of Zika virus RNA in semen for six months after returning from Haiti to Italy, January 2016. Eurosurveillance [Internet]. 2016 Aug 11 [cited 2016 Nov 17];21(32). Available from: http://www.eurosurveillance.org/ViewArticle.aspx?ArticleId=22556

11. Barzon L, Percivalle E, Pacenti M, Rovida F, Zavattoni M, Del Bravo P, et al. Virus and Antibody Dynamics in Travelers With Acute Zika Virus Infection. Clinical Infectious Diseases. 2018 Apr 3;66(8):1173–80.

12. Calvet GA, Kara EO, Giozza SP, Bôtto-Menezes CHA, Gaillard P, de Oliveira Franca RF, et al. Study on the persistence of Zika virus (ZIKV) in body fluids of patients with ZIKV infection in Brazil. BMC Infect Dis [Internet]. 2018 Jan 22 [cited 2019 Jan 7];18. Available from: https://www.ncbi.nlm.nih.gov/pmc/articles/PMC5778641/

13. Chung DH, Jonsson CB, Tower NA, Chu YK, Sahin E, Golden JE, et al. Discovery of a novel compound with anti-venezuelan equine encephalitis virus activity that targets the nonstructural protein 2. PLOS Pathogens. 2014 June;(6):e1004213.

14. Chung DH, Golden JE, Adcock RS, Schroeder CE, Chu YK, Sotsky JB, et al. Discovery of a Broad-Spectrum Antiviral Compound That Inhibits Pyrimidine Biosynthesis and Establishes a Type 1 Interferon-Independent Antiviral State. Antimicrob Agents Chemother. 2016 Aug 1;60(8):4552–62.

15. Severson WE, Shindo N, Sosa M, Fletcher T, White EL, Ananthan S, et al. Development and Validation of a High-Throughput Screen for Inhibitors of SARS CoV and Its Application in Screening of a 100,000-Compound Library. J Biomol Screen. 2007 Feb;12(1):33–40.

16. Kaptein SJF, Goethals O, Kiemel D, Marchand A, Kesteleyn B, Bonfanti JF, et al. A pan-serotype dengue virus inhibitor targeting the NS3-NS4B interaction. Nature. 2021 Oct;598(7881):504–9.

17. Kiemel D, Kroell ASH, Denolly S, Haselmann U, Bonfanti JF, Andres JI, et al. Pan-serotype dengue virus inhibitor JNJ-A07 targets NS4A-2K-NS4B interaction with NS2B/NS3 and blocks replication organelle formation. Nat Commun. 2024 July 19;15(1):6080.

18. Guo F, Wu S, Julander J, Ma J, Zhang X, Kulp J, et al. A Novel Benzodiazepine Compound Inhibits Yellow Fever Virus Infection by Specifically Targeting NS4B Protein. Diamond MS, editor. J Virol. 2016 Dec;90(23):10774–88.

19. Moquin SA, Simon O, Karuna R, Lakshminarayana SB, Yokokawa F, Wang F, et al. NITD-688, a pan-serotype inhibitor of the dengue virus NS4B protein, shows favorable pharmacokinetics and efficacy in preclinical animal models. Sci Transl Med. 2021 Feb 3;13(579):eabb2181.

20. Wang QY, Dong H, Zou B, Karuna R, Wan KF, Zou J, et al. Discovery of Dengue Virus NS4B Inhibitors. Journal of Virology. 2015 July 21;89(16):8233–44.

21. Zou J, Lee LT, Wang QY, Xie X, Lu S, Yau YH, et al. Mapping the Interactions between the NS4B and NS3 proteins of dengue virus. J Virol. 2015 Apr;89(7):3471–83.

22. Deng YQ, Zhang NN, Li CF, Tian M, Hao JN, Xie XP, et al. Adenosine Analog NITD008 Is a Potent Inhibitor of Zika Virus. Open Forum Infect Dis. 2016 Oct;3(4):ofw175.

23. Adcock RS, Chu YK, Golden JE, Chung DH. Evaluation of anti-Zika virus activities of broad-spectrum antivirals and NIH clinical collection compounds using a cell-based, high-throughput screen assay. Antiviral Res. 2017 Feb;138:47–56.

24. Hoffmann HH, Kunz A, Simon VA, Palese P, Shaw ML. Broad-spectrum antiviral that interferes with de novo pyrimidine biosynthesis. Proceedings of the National Academy of Sciences of the United States of America. 2011 Apr 5;(14):5777–82.

25. Lucas-Hourani M, Dauzonne D, Jorda P, Cousin G, Lupan A, Helynck O, et al. Inhibition of Pyrimidine Biosynthesis Pathway Suppresses Viral Growth through Innate Immunity. PLOS Pathog. 2013 Oct 3;9(10):e1003678.

26. Wang QY, Bushell S, Qing M, Xu HY, Bonavia A, Nunes S, et al. Inhibition of dengue virus through suppression of host pyrimidine biosynthesis. Journal of virology. 2011 July;(13):6548–56.

27. Luthra P, Naidoo J, Pietzsch CA, De S, Khadka S, Anantpadma M, et al. Inhibiting pyrimidine biosynthesis impairs Ebola virus replication through depletion of nucleoside pools and activation of innate immune responses. Antiviral Res. 2018 Oct;158:288–302.

28. Lipinski CA. Drug-like properties and the causes of poor solubility and poor permeability. Journal of Pharmacological and Toxicological Methods. 2000 July;44(1):235–49.

29. Veber DF, Johnson SR, Cheng HY, Smith BR, Ward KW, Kopple KD. Molecular properties that influence the oral bioavailability of drug candidates. J Med Chem. 2002 June 6;45(12):2615–23.

30. Doak BC, Kihlberg J. Drug discovery beyond the rule of 5 - Opportunities and challenges. Expert Opin Drug Discov. 2017 Feb;12(2):115–9.

31. Kaufusi PH, Tseng AC, Kelley JF, Nerurkar VR. Selective Reactivity of Anti-Japanese Encephalitis Virus NS4B Antibody Towards Different Flaviviruses. Viruses. 2020 Feb 14;12(2):212.

32. Miller S, Sparacio S, Bartenschlager R. Subcellular Localization and Membrane Topology of the Dengue Virus Type 2 Non-structural Protein 4B. Journal of Biological Chemistry. 2006 Mar;281(13):8854–63.

33. Li Y, Wong YL, Lee MY, Li Q, Wang Q, Lescar J, et al. Secondary Structure and Membrane Topology of the Full-Length Dengue Virus NS4B in Micelles. Angew Chem Int Ed. 2016 Sept 19;55(39):12068–72.

34. Kesteleyn B, Bonfanti JF, Bardiot D, De Boeck B, Goethals O, Kaptein SJF, et al. Discovery of JNJ-1802, a First-in-Class Pan-Serotype Dengue Virus NS4B Inhibitor. J Med Chem. 2024 Mar 14;67(5):4063–82.

35. Malyutina A, Majumder MM, Wang W, Pessia A, Heckman CA, Tang J. Drug combination sensitivity scoring facilitates the discovery of synergistic and efficacious drug combinations in cancer. Gallo J, editor. PLoS Comput Biol. 2019 May 20;15(5):e1006752.

36. Zheng S, Wang W, Aldahdooh J, Malyutina A, Shadbahr T, Tanoli Z, et al. SynergyFinder Plus: Toward Better Interpretation and Annotation of Drug Combination Screening Datasets. Genomics, Proteomics & Bioinformatics. 2022 June 1;20(3):587–96.

37. Roth NM, Reynolds MR, Lewis EL, Woodworth KR, Godfred-Cato S, Delaney A, et al. Zika-Associated Birth Defects Reported in Pregnancies with Laboratory Evidence of Confirmed or Possible Zika Virus Infection — U.S. Zika Pregnancy and Infant Registry, December 1, 2015–March 31, 2018. MMWR Morb Mortal Wkly Rep. 2022 Jan 21;71(3):73–9.

38. Bhatnagar J, Rabeneck DB, Martines RB, Reagan-Steiner S, Ermias Y, Estetter LBC, et al. Zika Virus RNA Replication and Persistence in Brain and Placental Tissue. Emerg Infect Dis. 2017 Mar;23(3):405–14.

39. Driggers RW, Ho CY, Korhonen EM, Kuivanen S, Jääskeläinen AJ, Smura T, et al. Zika Virus Infection with Prolonged Maternal Viremia and Fetal Brain Abnormalities. N Engl J Med. 2016 June 2;374(22):2142–51.

40. Sapparapu G, Fernandez E, Kose N, Cao B, Fox JM, Bombardi RG, et al. Neutralizing human antibodies prevent Zika virus replication and fetal disease in mice. Nature. 2016 Dec 15;540(7633):443–7.

41. Dunn MC, Knight DA, Waldman WJ. Inhibition of respiratory syncytial virus in vitro and in vivo by the immunosuppressive agent leflunomide. Antiviral therapy. 2011;(3):309–17.

42. El Kouni MH, Cha S. Isolation and partial characterization of a 5’-nucleotidase specific for orotidine-5’-monophosphate. Proc Natl Acad Sci U S A. 1982 Feb;79(4):1037–41.

43. Zheng Y, Li S, Song K, Ye J, Li W, Zhong Y, et al. A Broad Antiviral Strategy: Inhibitors of Human DHODH Pave the Way for Host-Targeting Antivirals against Emerging and Re-Emerging Viruses. Viruses. 2022 Apr 28;14(5):928.

44. Kaptein SJF, Goethals O, Kiemel D, Marchand A, Kesteleyn B, Bonfanti JF, et al. A pan-serotype dengue virus inhibitor targeting the NS3–NS4B interaction. Nature. 2021 Oct 21;598(7881):504–9.

45. Xie X, Wang QY, Xu HY, Qing M, Kramer L, Yuan Z, et al. Inhibition of Dengue Virus by Targeting Viral NS4B Protein. J Virol. 2011 Nov;85(21):11183–95.

46. Wang QY, Dong H, Zou B, Karuna R, Wan KF, Zou J, et al. Discovery of Dengue Virus NS4B Inhibitors. Diamond MS, editor. J Virol. 2015 Aug 15;89(16):8233–44.

47. Zou J, Xie X, Lee LT, Chandrasekaran R, Reynaud A, Yap L, et al. Dimerization of Flavivirus NS4B Protein. García-Sastre A, editor. J Virol. 2014 Mar 15;88(6):3379–91.

48. Varasteh Moradi S, Gagoski D, Mureev S, Walden P, McMahon KA, Parton RG, et al. Mapping Interactions among Cell-Free Expressed Zika Virus Proteins. J Proteome Res. 2020 Apr 3;19(4):1522–32.

49. Surya W, Tan P, Honey SS, Mehta D, Torres J. Flavivirus NS4B proteins do not form homodimers: discrepancies with an AlphaFold-based oligomeric model. Computational and Structural Biotechnology Journal. 2025;27:1660–72.

50. Płaszczyca A, Scaturro P, Neufeldt CJ, Cortese M, Cerikan B, Ferla S, et al. A novel interaction between dengue virus nonstructural protein 1 and the NS4A-2K-4B precursor is required for viral RNA replication but not for formation of the membranous replication organelle. PLoS Pathog. 2019 May;15(5):e1007736.

51. Li XD, Deng CL, Ye HQ, Zhang HL, Zhang QY, Chen DD, et al. Transmembrane Domains of NS2B Contribute to both Viral RNA Replication and Particle Formation in Japanese Encephalitis Virus. Perlman S, editor. J Virol. 2016 June 15;90(12):5735–49.

52. Zou J, Lee LT, Wang QY, Xie X, Lu S, Yau YH, et al. Mapping the Interactions between the NS4B and NS3 Proteins of Dengue Virus. Dermody TS, editor. J Virol. 2015 Apr;89(7):3471–83.

53. Umareddy I, Chao A, Sampath A, Gu F, Vasudevan SG. Dengue virus NS4B interacts with NS3 and dissociates it from single-stranded RNA. Journal of General Virology. 2006 Sept 1;87(9):2605–14.

54. Lu H, Zhan Y, Li X, Bai X, Yuan F, Ma L, et al. Novel insights into the function of an N-terminal region of DENV2 NS4B for the optimal helicase activity of NS3. Virus Research. 2021 Apr;295:198318.

55. Zou J, Xie X, Wang QY, Dong H, Lee MY, Kang C, et al. Characterization of Dengue Virus NS4A and NS4B Protein Interaction. Dermody TS, editor. J Virol. 2015 Apr;89(7):3455–70.

56. Muñoz-Jordán JL, Laurent-Rolle M, Ashour J, Martínez-Sobrido L, Ashok M, Lipkin WI, et al. Inhibition of Alpha/Beta Interferon Signaling by the NS4B Protein of Flaviviruses. J Virol. 2005 July;79(13):8004–13.

57. Wang Y, Xie X, Shi PY. Flavivirus NS4B protein: Structure, function, and antiviral discovery. Antiviral Res. 2022 Nov;207:105423.

58. Kiemel D, Kroell ASH, Denolly S, Haselmann U, Bonfanti JF, Andres JI, et al. Pan-serotype dengue virus inhibitor JNJ-A07 targets NS4A-2K-NS4B interaction with NS2B/NS3 and blocks replication organelle formation. Nat Commun. 2024 July 19;15(1):6080.

59. Porter SS, Gilchrist TM, Schrodel S, Tai AW. Dengue and Zika virus NS4B proteins differ in topology and in determinants of ER membrane protein complex dependency. J Virol. 2025 Feb 25;99(2):e0144324.

60. Ngo AM, Shurtleff MJ, Popova KD, Kulsuptrakul J, Weissman JS, Puschnik AS. The ER membrane protein complex is required to ensure correct topology and stable expression of flavivirus polyproteins. Elife. 2019 Sept 13;8:e48469.

61. Kim J, Alejandro B, Hetman M, Hattab EM, Joiner J, Schroten H, et al. Zika virus infects pericytes in the choroid plexus and enters the central nervous system through the blood-cerebrospinal fluid barrier. Pierson TC, editor. PLoS Pathog. 2020 May 1;16(5):e1008204.

62. Quick J, Grubaugh ND, Pullan ST, Claro IM, Smith AD, Gangavarapu K, et al. Multiplex PCR method for MinION and Illumina sequencing of Zika and other virus genomes directly from clinical samples. Nat Protoc. 2017 June;12(6):1261–76.

63. Li H. Minimap2: pairwise alignment for nucleotide sequences. Bioinformatics. 2018 Sept 15;34(18):3094– 100.

64. Li H, Handsaker B, Wysoker A, Fennell T, Ruan J, Homer N, et al. The Sequence Alignment/Map format and SAMtools. Bioinformatics. 2009 Aug 15;25(16):2078–9.

65. Oliinyk OS, Baloban M, Clark CL, Carey E, Pletnev S, Nimmerjahn A, et al. Single-domain near-infrared protein provides a scaffold for antigen-dependent fluorescent nanobodies. Nat Methods. 2022 June;19(6):740–50.

66. Li Y, Loh YR, Li Q, Luo D, Kang C. 1H, 15N and 13C backbone resonance assignment of the N-terminal region of Zika virus NS4B protein in detergent micelles. Biomol NMR Assign. 2025 June;19(1):1–6.

67. Castro MA, Parson KF, Beg I, Wilkinson MC, Nurmakova K, Levesque I, et al. Verteporfin is a substrate-selective γ-secretase inhibitor that binds the amyloid precursor protein transmembrane domain. Journal of Biological Chemistry. 2022 Apr;298(4):101792.

